# Phylogenomics reveals distinct evolutionary lineages of Japanese *Cinnamomum* species in the Nansei Islands

**DOI:** 10.64898/2026.09.23.753985

**Authors:** Sayaka Iizasa, Naoko Anjiki, Ei’ichi Iizasa

**Author notes:** Corresponding authors: Sayaka Iizasa Faculty of Agriculture, Kagoshima University 1-21-24 Korimoto, Kagoshima 890-0065, Japan Ei’ichi Iizasa Division of Psychosomatic Internal Medicine, Department of Social and Behavioral Medicine, Graduate School of Medical and Dental Sciences, Kagoshima University 8-35-1 Sakuragaoka, Kagoshima 890-8544, Japan.

## Abstract

Although *Cinnamomum* includes globally important spice and medicinal plants, genome-scale knowledge of *Cinnamomum* species native to Japan remains limited. The Nansei Islands are an approximately 1,200-km-long archipelago located between the Pacific Ocean and the East China Sea and represent a key biogeographic region shaped by deep-sea barriers such as the Tokara and Kerama Gaps. Here, we integrated chloroplast and nuclear genome data generated in this study with public datasets to examine the phylogenetic relationships of four *Cinnamomum* species native to the Nansei Islands and ten related species. Chloroplast genomes showed no major structural differences among the four island species, whereas nuclear genome data resolved two distinct evolutionary lineages in the Nansei Islands: *C. yabunikkei*, *C. × durifruticeticola*, and *C. daphnoides* formed a distinct Nansei Islands lineage separate from Sri Lankan, Indonesian, and Taiwanese lineages, whereas C. *sieboldii* was placed in a Taiwanese lineage. Ancestry estimation indicated that *C. × durifruticeticola* and *C. daphnoides* share a common ancestral component, while *C. sieboldii* exhibited a single homogeneous ancestral component, suggesting reduced ancestral diversity after divergence from a Taiwanese lineage. These findings reveal previously unrecognized genomic differentiation among Japanese *Cinnamomum* species and suggest that deep-sea barriers, dispersal, and long-term isolation shaped their diversification.

## Introduction

Species of the genus *Cinnamomum*, commonly known as cinnamon and cassia, include globally important spice and medicinal plants that have been used since ancient times. Owing to their scientific and commercial value, numerous morphological and genetic studies have been conducted on *Cinnamomum* species across Asia and surrounding regions^1–4^. However, knowledge of *Cinnamomum* species endemic to the Nansei Islands, Japan, remains limited.

The Nansei Islands comprise an approximately 1,200-km-long archipelago extending from the southern tip of Kyushu (the southwesternmost of Japan’s four main islands) to Taiwan, where several species of *Cinnamomum* occur naturally (Fig. 1a). Within this archipelago, the Tokara Gap and the Kerama Gap are located in its northern and southern regions, respectively (Fig. 1a)^5^. These deep marine gaps are thought to have remained oceanic barriers throughout the Pliocene to the Pleistocene, and numerous endemic species occur on the islands between them^5,6^. *Cinnamomum sieboldii* Meisn., commonly known as Nikkei or Japanese cinnamon, is an evergreen tree species endemic to the Nansei Islands and is classified as a near-threatened tree species in Japan. This species has been used for various culinary and medicinal applications for many centuries, however, scientific investigations into its properties remain limited. Previous studies have characterized the isolation of bark extracts from *C. sieboldii* and elucidated aspects of its phytochemical composition^7^. In addition, although comparative anatomical studies examining bark and root bark structures among several Japanese cinnamon species, including *C. sieboldii*, have been reported^8^, phylogenetic relationships with other recognized species within the genus *Cinnamomum* remain unresolved.

**Fig. 1.**
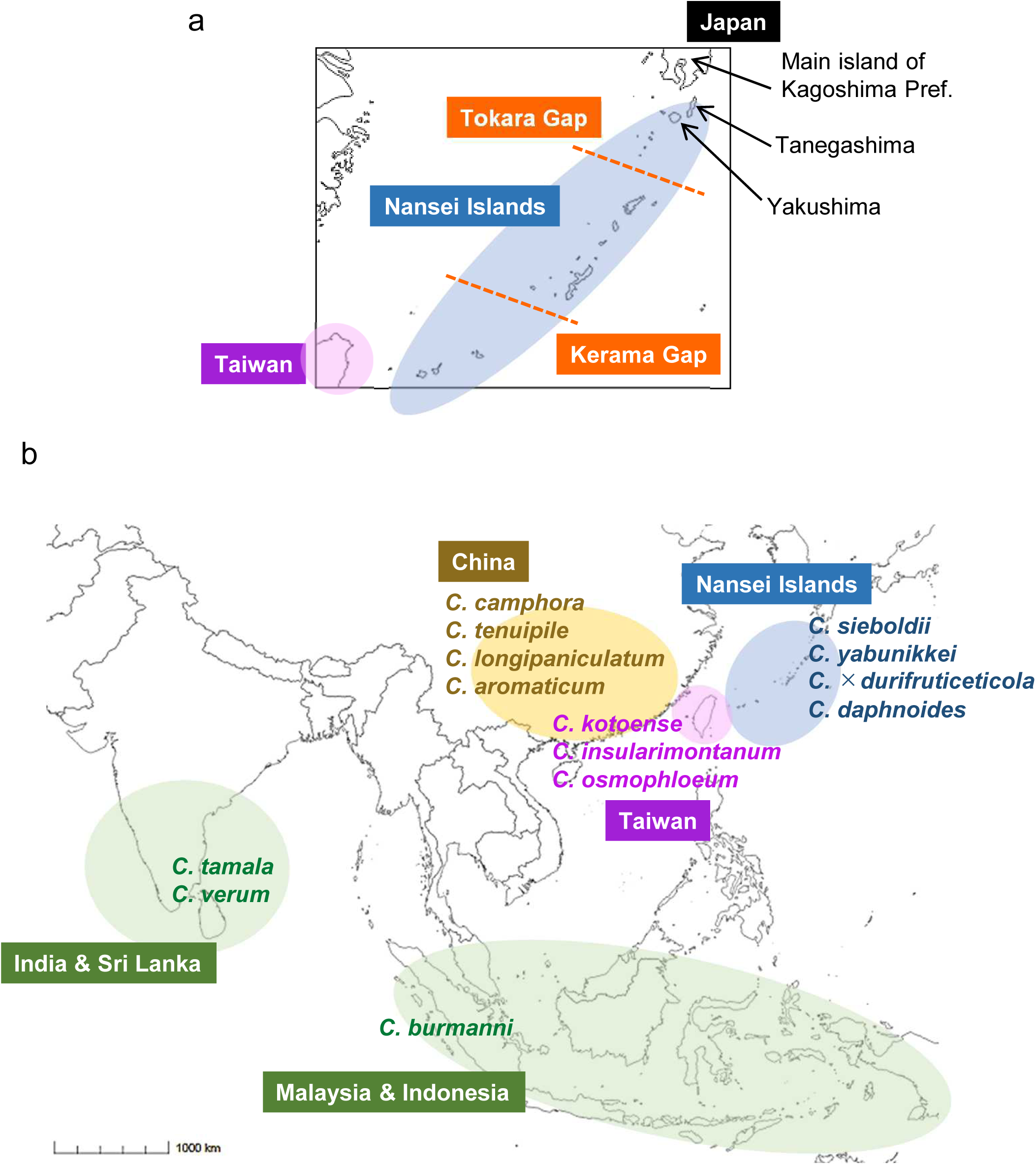
(a) The enlarged map shows the distribution of islands across the Nansei Islands. Orange dotted lines indicate the positions of two major gaps. Sampling sites are indicated. (b) The main habitats of the *Cinnamomum* taxa plants used in this study.

Not only *C. sieboldii*, but also *C. yabunikkei* H.Ohba, *C. daphnoides* Siebold et Zucc., and *C. × durifruticeticola* Hatus. occur in the Nansei Islands (Fig. 1b). *C. yabunikkei* H.Ohba (syn. *C. japonicum* Siebold ex Nakai), is an evergreen tree native to coastal forests of East Asia, including Korea, China, and Japan. This species has long been the subject of taxonomic debate, particularly regarding whether *C. chenii* and *C. chekiangense* represent distinct species. However, recent phylogenetic analyses by Lin *et al*. (2023) demonstrated that specimens identified as *C. chenii* are nested within the *C. yabunikkei* clade, thereby supporting the conclusion that these taxa are conspecific^9^. *C. daphnoides* Siebold et Zucc. is a rare, near-threatened tree species in Japan, known for its elegant form, lustrous foliage, and adaptations such as wind resistance, drought tolerance, salt spray tolerance, and high light endurance^10^. *C. daphnoides* has generally been regarded as a Japanese endemic species, although a report has documented its occurrence on coastal cliffs in Xiangshan County, Zhejiang Province, China^11^. *C. × durifruticeticola* Hatus. is believed to be a hybrid taxon derived from a cross between *C. yabunikkei* and *C. daphnoides*^12^. This species has been classified as a rare species in local flora surveys. As phylogenetic relationships within the genus *Cinnamomum* have been increasingly clarified in recent years, a focused investigation of Japanese *Cinnamomum* species inhabiting the Nansei Islands, including *C. sieboldii*, by a comprehensive genomic analysis will further refine our understanding of their evolutionary history and provide new insights^1,2,13^.

The chloroplast (cp) genome is commonly employed to investigate evolutionary relationships among plants; however, because cpDNA represents a single lineage inherited from only one parent and evolves independently of the nuclear genome, phylogenetic inference based solely on cp genome sequences has important limitations. Recently, an approach has been reported in which the same raw read sequencing data are used to construct both cp genome–based phylogenetic trees and nuclear DNA phylogenies using Read2Tree, a cost-efficient method for extracting conserved nuclear gene sequences directly from raw read data^14,15^. In this study, we applied this approach to perform both cp genome–based and nuclear DNA–based phylogenetic analyses. In addition, population structure analyses were performed to further clarify phylogenetic relationships within the genus. This comprehensive approach allowed us to identify genomic differentiation among *Cinnamomum* lineages in the Nansei Islands, clarify their phylogenetic relationships with known lineages, and evaluate whether these genetic patterns align with currently recognized species boundaries

## Results

### Four *Cinnamomum* species of Nansei Islands and other recognized species within the genus

In this study, we examined four *Cinnamomum* species occurring in the Nansei Islands —*C. sieboldii*, *C. yabunikkei*, *C. × durifruticeticola*, and *C. daphnoides*—and compared them with other 10 taxonomically recognized species. These included *C. camphora*, *C. tenuipile*, *C. longipaniculatum*, and *C. aromaticum*, which are native to China; *C. kotoense*, *C. insularimontanum*, and *C. osmophloeum*, which are native to Taiwan; and *C. tamala*, *C. verum*, and *C. burmanni*, which are native to India, Sri Lanka, Malaysia, and Indonesia (Fig. 1b).

The morphological features of four *Cinnamomum* species of Nansei Islands were shown on Fig. 2. *C. sieboldii* and *C. yabunikkei* possess long, narrow leaves with acuminate apices, whereas *C. daphnoides* and *C. × durifruticeticola* exhibit relatively small, rounded leaves. In addition, naked-eye observations indicated that the adaxial leaf surfaces of *C. sieboldii* and *C. yabunikkei* are lustrous, whereas those of *C. daphnoides* and *C. × durifruticeticola* appear dull.

**Fig. 2.**
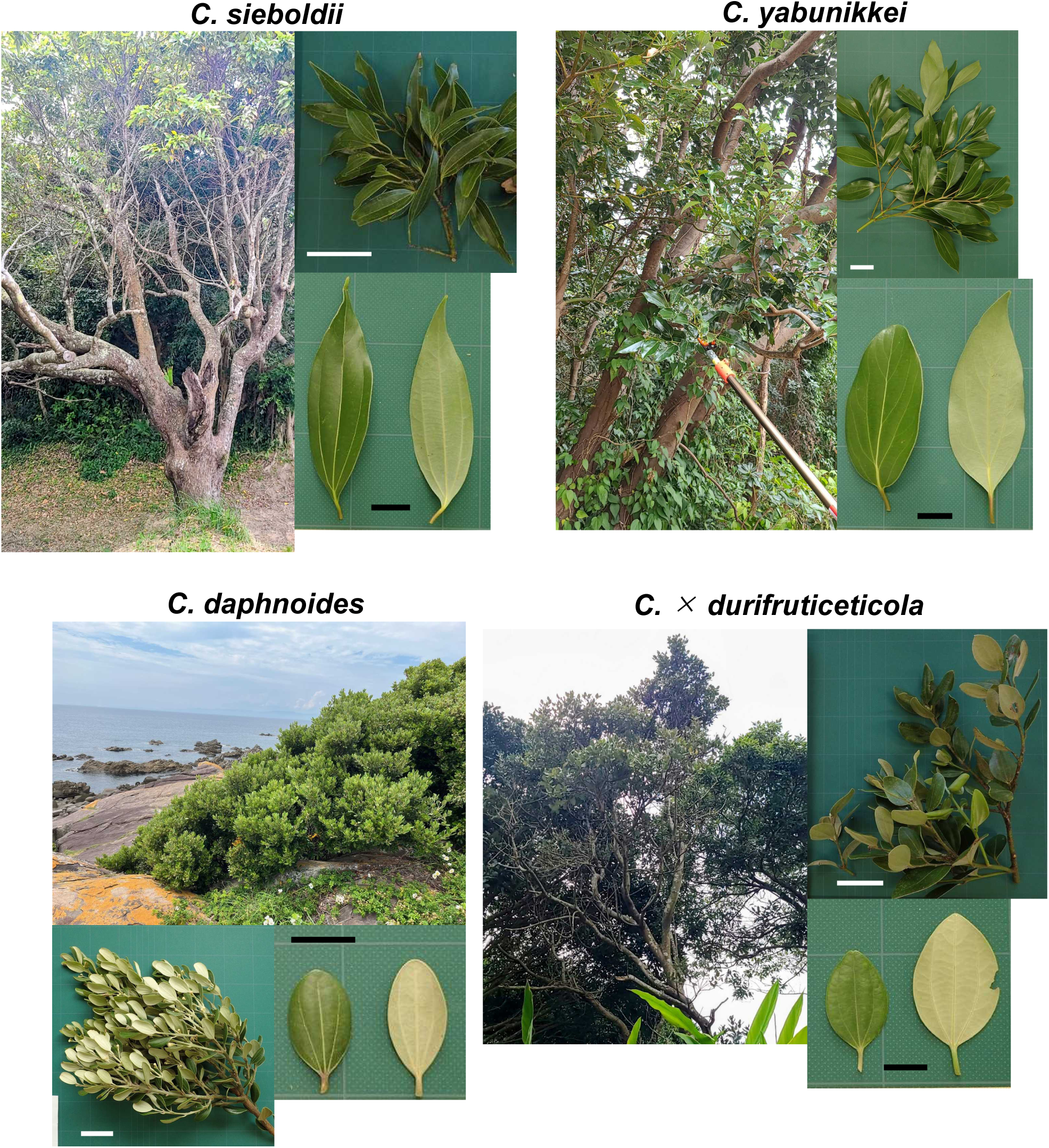
Morphological features of *C. sieboldii*, *C. yabunikkei*, *C. daphnoides*, and *C. × durifruticeticola*. Images show the whole plant, branchlets, and leaves. White scale bars = 5 cm; black scale bars = 2 cm.

### Basic characteristics of the chloroplast genomes of sampled *Cinnamomum* species

In the Nansei Islands, the Kerama Gap, a major biogeographical boundary, separates the biota of Taiwan and the southern Nansei Islands from that of the central and northern Nansei Islands (Fig. 1a)^16–18^. Accordingly, we collected plant samples of the genus *Cinnamomum* from the Nansei Islands located north of the Kerama Gap. Leaf samples of *C. sieboldii* and *C. × durifruticeticola* were obtained from individuals originally collected on Tanegashima Island and maintained at a stock center (Fig. 1a). *C. insularimontanum* and *C. verum,* both non-native *Cinnamomum* species, were also sampled from the same stock center. *C. verum* was included as a reference to validate the analytical results and the reliability of species identification. Leaves of *C. yabunikkei* were collected from Yakushima Island (30.2406° N, 130.2551° E) (Fig. 1a). For comparative purposes, one individual of *C. sieboldii* was collected from the main island of Kagoshima Prefecture, Kyushu (*C. sieboldii*_2; 31.5435° N, 130.2703° E) (Fig. 1a).

After quality control, 3.84–4.28 Gb of high-quality clean sequencing data were obtained. First, we assembled and annotated the cp genomes of these 6 samples and compared them to that of *C. daphnoides* obtained from a public database. The assembled cp genomes ranged from 152,762 to 154,176 bp in length and exhibited a typical quadripartite structure, consisting of a pair of inverted repeat (IR) regions (20,074–20,892 bp) separated by a large single-copy (LSC) region (93,685–93,721 bp) and a small single-copy (SSC) region (18,833–18,916 bp) (Fig. 3., Table 1, and Supplementary Information Fig. S1). The overall GC content of the complete cp genomes was 39.16–39.18% (Table 1). These structural features were highly conserved among the six samples of five analyzed species. All cp genomes encoded 126 functional genes, comprising 81 protein-coding genes, 37 transfer RNA (tRNA) genes, and 8 ribosomal RNA (rRNA) genes (Table 1). Among these, 15 genes contained a single intron (*ndhA, ndhB, petB, petD, atpF, rpl2, rpl16, rpoC1, rps16, trnA-UGC, trnG-UCC, trnI-GAU, trnL-UAA, trnK-UUU* and *trnV-UAC*), whereas two genes contained two introns (*pafI* and *clpP1*), with *rps12* undergoing trans-splicing. This gene and intron content was consistent across the analyzed species, indicating that the structure of the cp genome is conserved.

**Fig. 3.**
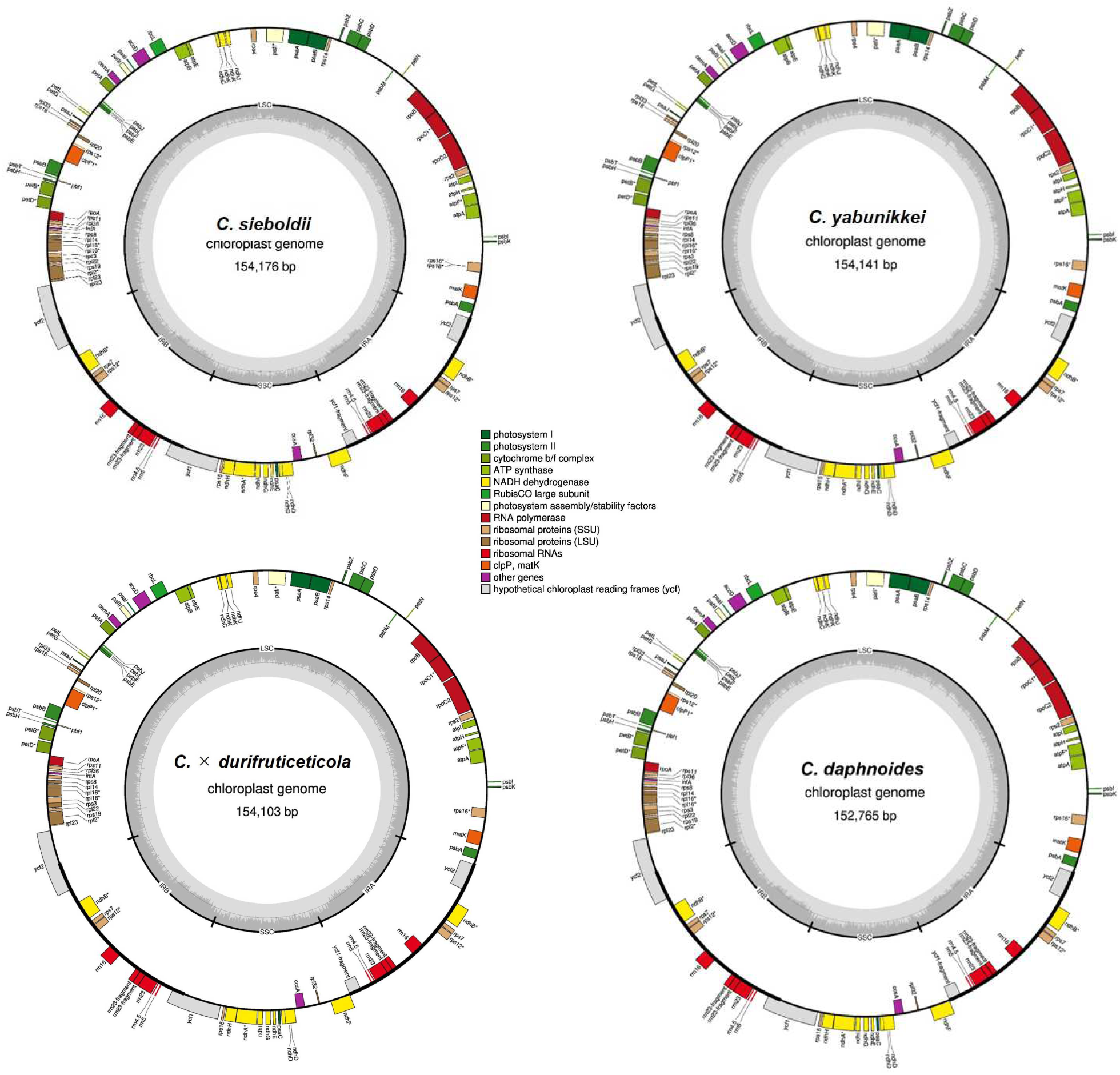
The complete plastome map of *C. sieboldii*, *C. yabunikkei*, *C. × durifruticeticola* and *C. daphnoides*^10^. Conserved plastid genes are shown as colored boxes, with transcription proceeding clockwise on the inner circle and counterclockwise on the outer circle. Gray bars in the central ring depict the plastome GC content.

**Table 1.**
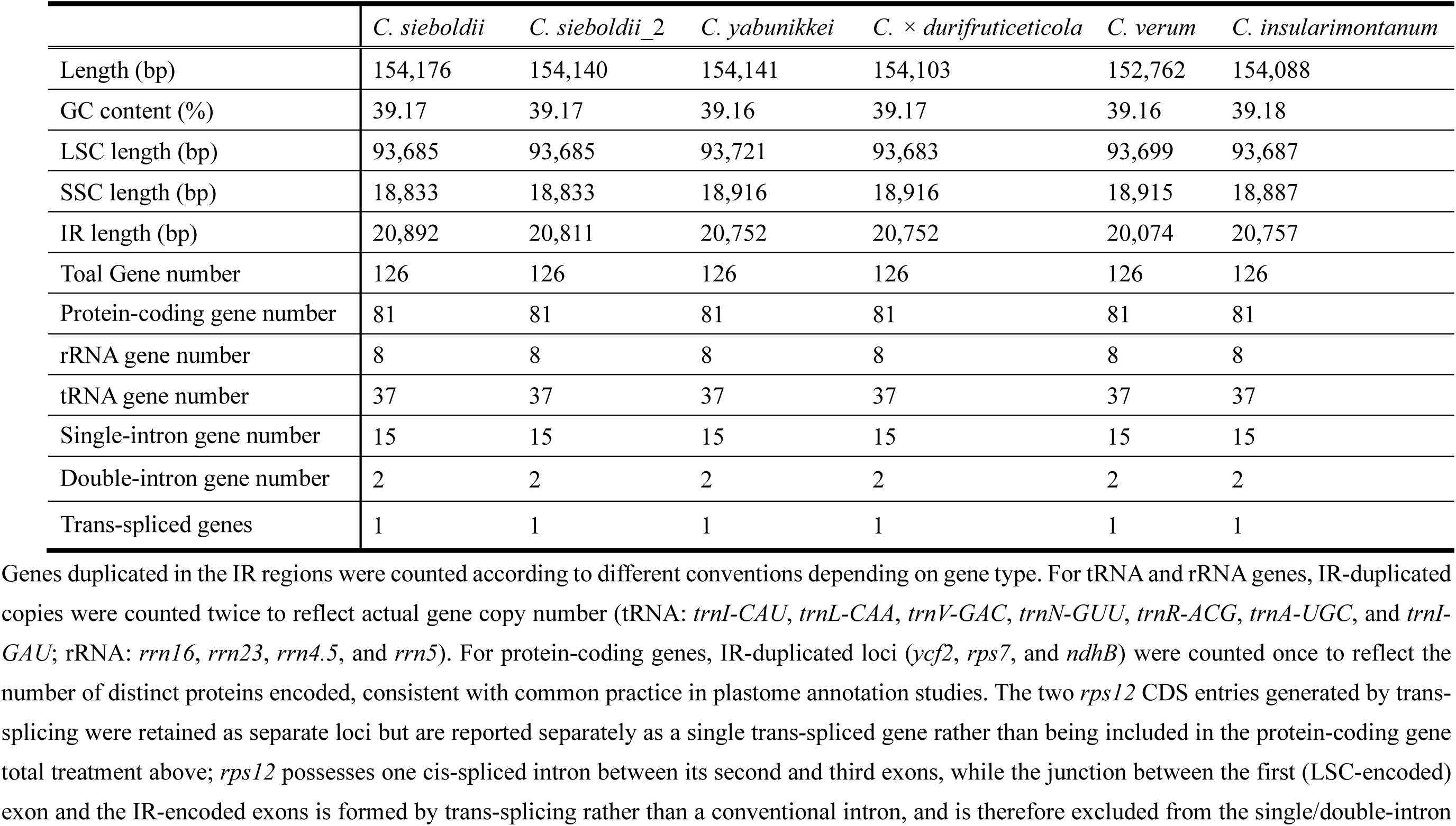
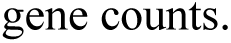
Comparison of the basic characteristics of the cp genomes of six *Cinnamomum* species we analyzed.

|  | *C. sieboldii* | *C. sieboldii_2* | *C. yabunikkei* | *C. × durifruticeticola* | *C. verum* | *C. insularimontanum* |
| --- | --- | --- | --- | --- | --- | --- |
| Length (bp) | 154,176 | 154,140 | 154,141 | 154,103 | 152,762 | 154,088 |
| GC content (%) | 39.17 | 39.17 | 39.16 | 39.17 | 39.16 | 39.18 |
| LSC length (bp) | 93,685 | 93,685 | 93,721 | 93,683 | 93,699 | 93,687 |
| SSC length (bp) | 18,833 | 18,833 | 18,916 | 18,916 | 18,915 | 18,887 |
| IR length (bp) | 20,892 | 20,811 | 20,752 | 20,752 | 20,074 | 20,757 |
| Toal Gene number | 126 | 126 | 126 | 126 | 126 | 126 |
| Protein-coding gene number | 81 | 81 | 81 | 81 | 81 | 81 |
| rRNA gene number | 8 | 8 | 8 | 8 | 8 | 8 |
| tRNA gene number | 37 | 37 | 37 | 37 | 37 | 37 |
| Single-intron gene number | 15 | 15 | 15 | 15 | 15 | 15 |
| Double-intron gene number | 2 | 2 | 2 | 2 | 2 | 2 |
| Trans-spliced genes | 1 | 1 | 1 | 1 | 1 | 1 |

Genes duplicated in the IR regions were counted according to different conventions depending on gene type. For tRNA and rRNA genes, IR-duplicated copies were counted twice to reflect actual gene copy number (tRNA: *trnI-CAU*, *trnL-CAA*, *trnV-GAC*, *trnN-GUU*, *trnR-ACG*, *trnA-UGC*, and *trnI-GAU*; rRNA: *rrn16*, *rrn23*, *rrn4.5*, and *rrn5*). For protein-coding genes, IR-duplicated loci (*ycf2*, *rps7*, and *ndhB*) were counted once to reflect the number of distinct proteins encoded, consistent with common practice in plastome annotation studies. The two *rps12* CDS entries generated by trans-splicing were retained as separate loci but are reported separately as a single trans-spliced gene rather than being included in the protein-coding gene total treatment above; *rps12* possesses one cis-spliced intron between its second and third exons, while the junction between the first (LSC-encoded) exon and the IR-encoded exons is formed by trans-splicing rather than a conventional intron, and is therefore excluded from the single/double-intron
gene counts.

### The cp genome-based phylogenetic relationship of *Cinnamomum* species of Nansei Islands and related taxa

To investigate evolutionary relationships and maternal lineage patterns, we constructed a chloroplast (cp) genome–based phylogenetic tree using 17 *Cinnamomum* species. Complete cp genome sequences for species other than *C. sieboldii*, *C. sieboldii_2*, *C. yabunikkei*, *C. × durifruticeticola*, *C. insularimontanum*, and *C. verum* were obtained from public databases (Supplementary Information Table S1). In addition, for verification purposes, an independently assembled cp genome of *C. verum* from a public database (*C. verum_2*) was included. Two species—*C. micranthum* and *C. foveolatum*—were designated as outgroup taxa.

Phylogenetic analyses based on the maximum likelihood (ML) method resolved four well-supported clades (Fig. 4). Notably, all four *Cinnamomum* species of Nansei Islands were clustered within clade IV, together with *C. osmophloeum* and *C. insularimontanum*. This clustering reflects shared maternal evolutionary histories and suggests a common matrilineal origin. Within clade IV, *C. sieboldii* formed a small group that is sister related to *C. insularimontanum,* whereas *C. yabunikkei*, *C. × durifruticeticola*, and *C. daphnoides* also formed a small group that is sister related to *C. osmophloeum,* supporting their close phylogenetic relationships. Overall, it appears that following the early divergence of the Sri Lankan lineage, which includes *C. verum*, clades comprising the Chinese lineage, the Indonesian lineage, and subsequently the Taiwanese and Nansei Islands lineages diverged in succession.

**Fig. 4.**
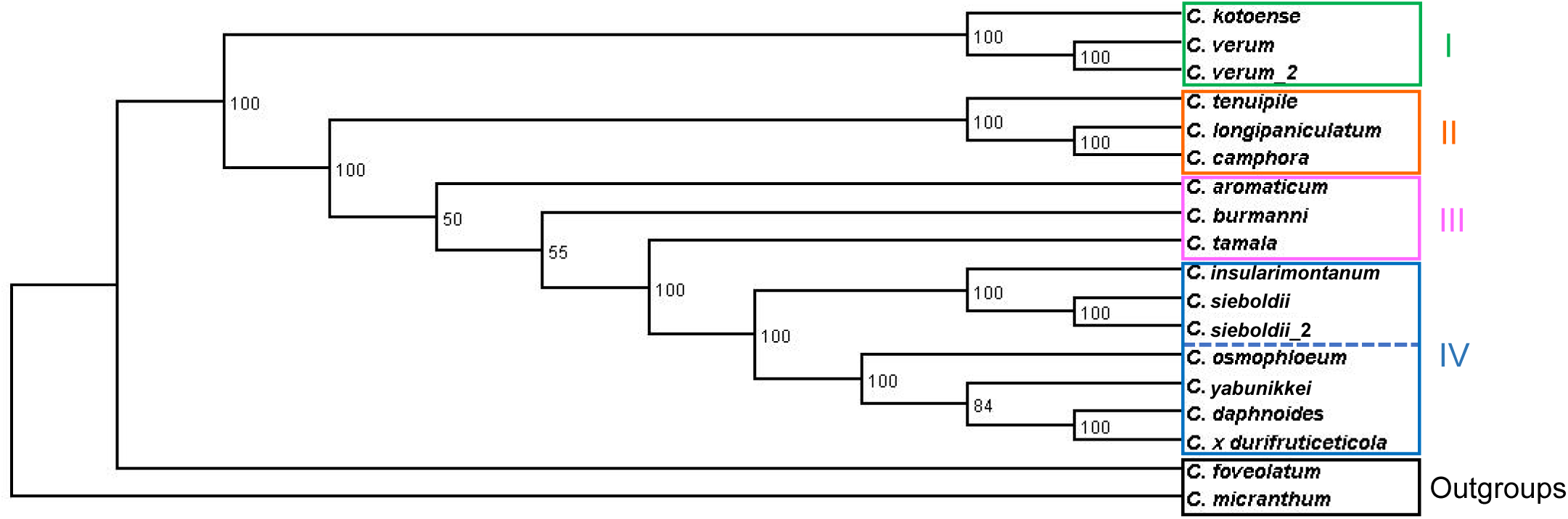
Maximum likelihood (ML) phylogenetic cp tree of 16 *Cinnamomum* taxa. The accession ID is provided in Supplementary Information Table S1 and S2. Nodal values represent ML bootstrap values. The tree is shown as rerooted for visualization purposes; no evolutionary direction is implied. *C. micranthum* and *C. foveolatum* were designated as outgroup taxa

### The nuclear genome-based phylogenetic relationship of *Cinnamomum* species of Nansei Islands and related taxa

Using nuclear DNA, which is characterized by parental inheritance and a high mutation rate, we conducted a phylogenetic analysis that reflects more comprehensive genetic information. Using the same reads from which the cp genome data was obtained, we used Read2Tree to retrieve sequences from various conserved nuclear genes and generate an ML tree. A tree was constructed based on 14 species. Nuclear genome sequences for species other than *C. sieboldii*, *C. sieboldii_2*, *C. yabunikkei*, *C. × durifruticeticola*, *C. insularimontanum*, and *C. verum* were obtained from public databases (Supplementary Information Table S1).

The generated phylogenetic tree yielded five distinct groups (Fig. 5). Notably, *C. sieboldii* was clustered within clade_4, together with *C. osmophloeum* and *C. insularimontanum*, whereases *C. yabunikkei*, *C. × durifruticeticola*, and *C. dahpnoides* were clustered within clade_5. Clade_5 was composed of these three species and did not include any others, indicating the unique phylogenetic relationship among them. This clade appears to have diverged after the split of the Sri Lankan lineage containing *C. verum*, but prior to the diversification of Indonesian and Taiwanese lineages. Clade_3 and _4 formed a large group consisting of seven species of various regional origins, including *C. sieboldii*. Taiwanese and Nansei Islands species were closely related or intermingled in the cp phylogeny, whereas the nuclear phylogeny did not always recover the same relationships.

**Fig. 5.**
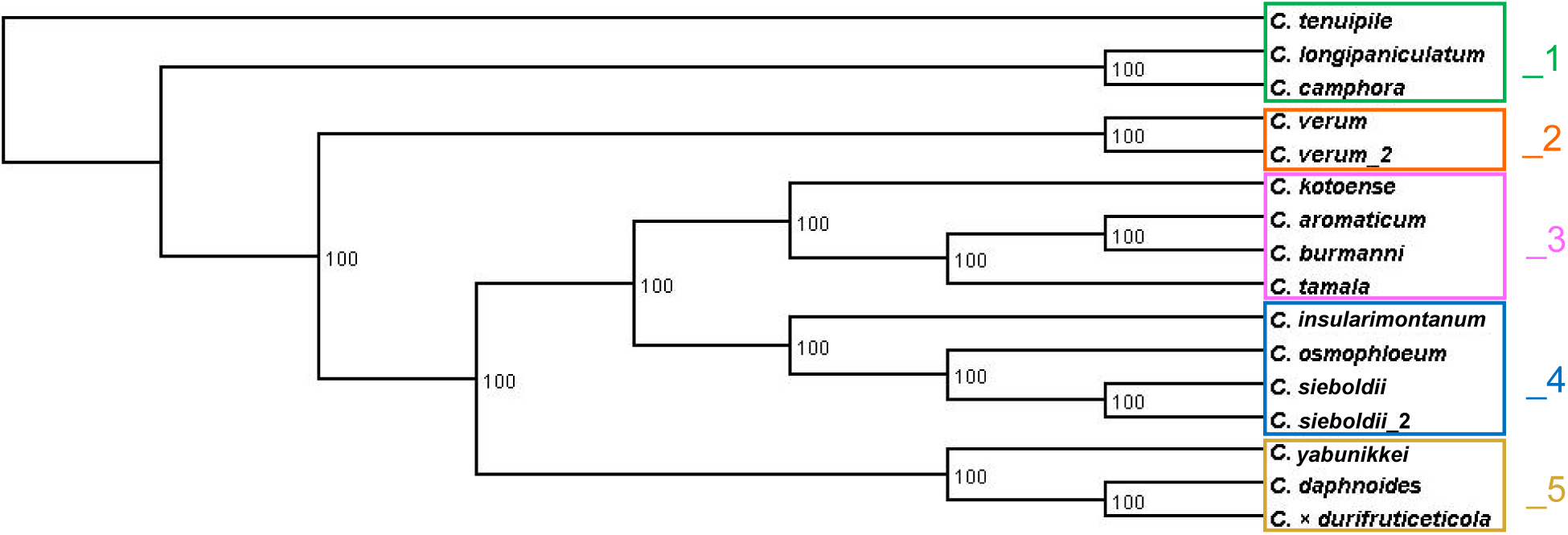
ML phylogenetic nuclear gene tree of 14 *Cinnamomum* taxa. The accession ID is provided in Supplementary Information Table S1 and S2. Nodal values represent ML bootstrap values. The tree is shown as rerooted for visualization purposes; no evolutionary direction is implied.

### Population structure visualized by principal component analysis and sparse non-negative matrix factorization

To visualize genome-wide genetic similarity and population structure among taxa, we first performed principal component analysis (PCA) based on the sequence data matrix. The first three principal components explained 33.2%, 13.6%, and 8.7% of the total variance, respectively. The PCA plot revealed five distinct clusters (Fig. 6a). Cluster_1 comprised *C. temuipile*, *C. longipaniculatum,* and *C. camphora*. Cluster_2 included two samples of *C. verum*. Cluster_3 consisted of *C. aromaticum*, *C. burmanni*, and *C. tamala*. Cluster_4 comprised *C. kotoense*, *C. osmophloeum*, *C. insularimontanum*, and two samples of *C. sieboldii*. Cluster_5 included *C. yabunikkei*, *C. × durifruticeticola*, and *C. daphnoides*. The clustering pattern observed in the PCA was largely congruent with the phylogenomic tree (Fig. 5), indicating that species-level divergence was supported by genome-wide genetic variation. Consistent with the phylogenetic results, *C. sieboldii* was separated from *C. yabunikkei*, *C. × durifruticeticola*, and *C. daphnoides*. In contrast, although *C. kotoense* was closely related to *C. aromaticum*, *C. burmanni*, and *C. tamala* in the phylogenetic analysis, PCA placed *C. kotoense* in a distinct cluster. Furthermore, cluster_4 was found to be genetically intermediate between the other clusters, suggesting possible genetic admixture.

**Fig. 6.**
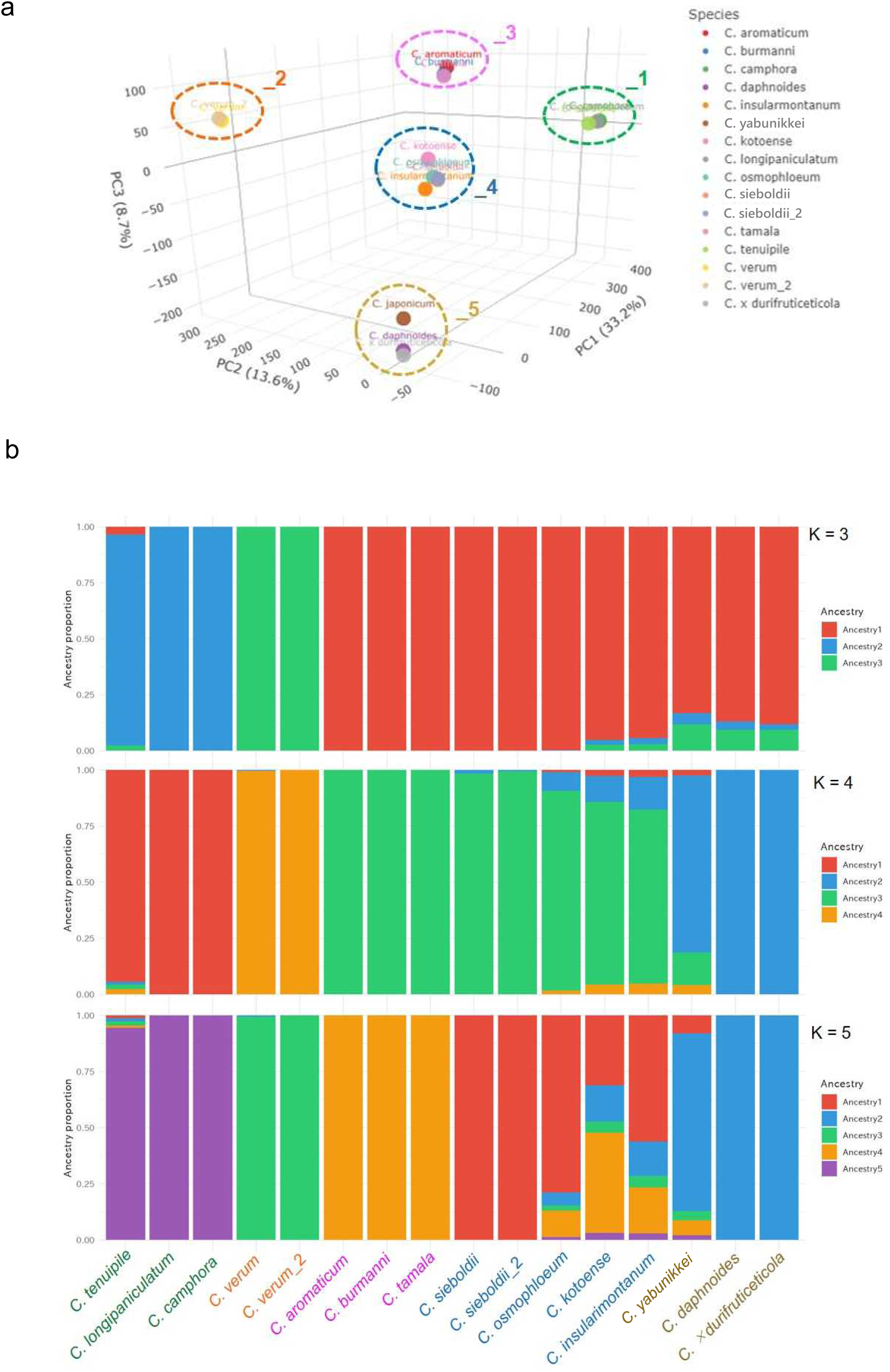
Population structure in the 14 *Cinnamomum* taxa individuals. (a) Three-dimensional plot of principal component analysis for 16 ingroup samples of *Cinnamomum*. Clusters were outlined and colored to correspond to the clades in the phylogenetic tree shown in Fig. 5. (b) Population structure analysis using sNMF. Barplot (K = 3, 4, 5) showing individual ancestry proportions, ordered according to their phylogenetic relationships. Taxa names were colored to correspond to the PCA clusters shown in Fig. 6a.

Ancestry estimation was further examined using sparse non-negative matrix factorization (sNMF). At K = 3 and K = 4, *C. sieboldii* clustered with *C. aromaticum*, *C. burmanni*, and *C. tamala*, suggesting that they shared a common ancestor (Fig. 6b). At K = 3, *C. yabunikkei*, *C. × durifruticeticola*, and *C. daphnoides* grouped together with *C. osmophloeum*, *C. kotoense*, and *C. insularimontanum*, showing partial admixture. At K = 4, *C. × durifruticeticola* and *C. daphnoides* formed a distinct population without admixture. At K = 5, the population structure was further resolved into five ancestral components, including one group comprising all five populations; *C. sieboldii* formed an independent population, whereas *C. yabunikkei* was assigned to the latter. Overall, these patterns support the presence of genetic admixture in cluster_4 identified by PCA. Consistent with this interpretation, the cross-entropy criterion of the sNMF analysis identified K = 5 as the most likely number of ancestral populations (Supplementary Information Fig. S2).

## Discussion

We constructed phylogenetic trees derived from the cp genome and the nuclear genome based on a single set of genomic data. In particular, we used the Read2Tree pipeline to infer a phylogenetic tree based on nuclear genomes^15^. The resulting nuclear phylogenetic tree was largely consistent with a previously published topology based on whole-genome resequencing data^13^, supporting the validity of the Read2Tree approach for nuclear phylogenomic inference in this genus. In contrast, the cp genome–based phylogeny showed partial discordance with the nuclear tree (Figs. 3 and 4). Such cytonuclear incongruence may reflect chloroplast capture following historical hybridization, incomplete lineage sorting (ILS), or the retention of ancestral maternal chloroplast lineages. In particular, the Taiwanese, Nansei Islands, and Indonesian lineages, which form clades III and IV in the cp-based phylogeny, may have experienced a complex evolutionary history involving maternal chloroplast introgression or the persistence of ancient cp lineages.

Comparative analyses of cp genomes revealed no significant differences in genomic structure among the four *Cinnamomum* species occurring in the Nansei Islands (Fig. 3). Nevertheless, phylogenetic analyses demonstrated that, *C. sieboldii* was clearly separated from the remaining three species—*C. yabunikkei*, *C. × durifruticeticola*, and *C. daphnoides* (Fig. 4 and Fig. 5). In the nuclear phylogenomic tree, *C. sieboldii* was most closely related to the Taiwanese species *C. osmophloeum* and shared common ancestry with a Taiwanese lineage (Fig. 5 and Fig. 6b). This finding indicates that, despite its current distribution in the Nansei Islands, the nuclear genome of *C. sieboldii* retains a strong affinity with Taiwanese lineages, suggesting that the species likely originated from a dispersal event from Taiwan to the Nansei Islands. Furthermore, the nuclear phylogeny places the divergence of *C. sieboldii* after that of the Taiwanese species, implying a post-Miocene origin, given that several Taiwanese *Cinnamomum* species have been reported to have diverged during the Miocene^13^. This evolutionary scenario is consistent with geological evidence indicating that crustal thinning in the western East China Sea from the late Miocene to the middle Pliocene caused eastward drift of the Ryukyu Arc, and that between approximately 1.6 and 1.3 Ma, much of the East China Sea region was subaerial, potentially connecting the Ryukyu Arc to the Eurasian continent^19^. It is therefore plausible that *C. sieboldii* diverged from a Taiwanese lineage during this period of geological transition. Following this dispersal, gene flow or hybridization among multiple lineages may have occurred more frequently in Taiwan, whereas gene flow from external lineages into the Nansei Islands was likely restricted by geographical barriers such as the Tokara and Kerama Gaps, together with inter-island isolation. As a result, *C. sieboldii* may exhibit reduced ancestral diversity, as reflected by the single ancestral component detected in the sNMF analysis (Fig. 6b). Because no molecular clock analyses were conducted in this study and divergence time estimates rely on previously published data^13^ these interpretations should be considered provisional.

Island environments promote distinctive patterns of genetic differentiation and evolutionary divergence by isolating terrestrial habitats through geological barriers, thereby facilitating the long-term preservation of biological diversity. In the Nansei Islands, the Kerama Gap, a major biogeographical boundary, separates the biota of Taiwan and the southern Nansei Islands from that of the central and northern Nansei Islands^16–18^. This Gap has been regarded as a strong geographical barrier for terrestrial animals^17,18^, its effect on plants may be more permeable. In some plant taxa, including fleshy-fruited genera such as *Cinnamomum*, long-distance dispersal across marine barriers can occur via bird-mediated seed dispersal, and potentially through other species-specific mechanisms. Consequently, the Kerama Gap may have functioned as a “filter barrier” rather than an absolute barrier for plants, selectively restricting gene flow depending on dispersal mode, dispersal frequency, and ecological suitability for establishment.

The evolutionary status of *C. daphnoides* and *C. × durifruticeticola* has long remained unclear. Although the complete cp genome of *C. daphnoides* has recently been reported^10^, its phylogenetic position within the genus *Cinnamomum*, as determined by the nuclear genome, remains poorly understood. Furthermore, *C. × durifruticeticola* has been hypothesized to represent a hybrid between *C. yabunikkei* and *C. daphnoides*, although this assumption has not been supported by genetic evidence. In the nuclear phylogenomic tree, *C. yabunikkei*, *C. daphnoides*, and *C. × durifruticeticola* formed a Nansei Islands clade that is distinct from Taiwanese clade (Fig. 5). Furthermore, sNMF analysis revealed that *C. × durifruticeticola* and *C. daphnoides* constitute a single genetic population derived from a common ancestor, indicating that these taxa form a cohesive evolutionary lineage rather than representing independently derived or recently admixed entities (Fig. 6b). This finding is particularly notable given the long-standing treatment of *C. × durifruticeticola* as a hybrid; instead, its genetic structure closely mirrors that of *C. daphnoides*, suggesting a shared and stable evolutionary history. Unlike *C. sieboldii*, which exhibits close genetic affinities with Taiwanese species, this lineage appears to have diverged earlier and evolved independently, consistent with a prolonged evolutionary history within the Nansei Islands. Notably, the presence of clade_5 in the nuclear genome phylogeny and cluster_5 in the PCA does not fully align with the cp genome phylogeny, implying that finer-scale nuclear differentiation exists within this group that is not captured by cp genome data alone. The placement of cluster_5 at the lower end of the PC1 axis further suggests the retention of relatively ancestral genomic features compared with other examined populations.

Interestingly, *C. yabunikkei* exhibited multiple ancestral components, one of which corresponds to the ancestry characteristic of *C. × durifruticeticola* and *C. daphnoides*. Although phylogenetic analyses indicate that *C. yabunikkei* diverged prior to these lineages, the sNMF results raise the possibility of secondary gene flow from the *C. × durifruticeticola*–*C. daphnoides* lineage into *C. yabunikkei* following divergence. Consequently, these findings do not support a simple hybrid-origin hypothesis for *C. × durifruticeticola*, but instead suggest that this lineage retained an independent genetic background while later experiencing genetic contact with *C. yabunikkei*. Nevertheless, a limitation of this study is that only a single individual was sampled for several species, potentially underrepresenting intraspecific genetic variation. Broader sampling across the distribution ranges of these taxa will be essential to further validate the evolutionary scenarios proposed here.

Taken together, these findings indicate that the evolutionary history of *Cinnamomum* in the Nansei Islands cannot be explained by a simple unidirectional dispersal from Taiwan. Rather, it is more accurately interpreted as reflecting a complex island evolutionary history shaped by the interplay of three factors: (1) the independent colonization of *C. sieboldii*, which is phylogenetically affiliated with the Taiwanese lineage; (2) the preservation of an ancient, independently diverged Nansei Islands lineage; and (3) the sharing or transfer of maternal cp genomes as evidenced in cp genome analyses.

Our findings advance our understanding of the evolutionary history of *Cinnamomum* species in the Nansei Islands and provide important insights into the genomic dynamics and interspecific relationships within this genus, highlighting the broader scientific significance of this archipelago as a potential center of plant diversification. Moreover, many *Cinnamomum* species inhabiting the Nansei Islands are classified as endangered or rare. Given the evolutionary distinctiveness of the Nansei Islands *Cinnamomum* lineages revealed in this study, we emphasize the urgent need for conservation measures to protect these irreplaceable components of the regional flora.

## Materials and Methods

### Sample collection, DNA extraction, and sequencing

*C. sieboldii*, *C. verum*, *C. insularimontanum*, and *C. × durifruticeticola* were collected from cultivated material at the Tanegashima Division, Reseach Center for Medicinal Plant Resources in Kagoshima Prefecture, Japan. *C. yabunikkei* and *C. sieboldii*_2 were sampled in Yakushima and main island of Kagoshima Prefecture, respectively. The origins of these plants are described in Fig. 1. All fresh leaves were transported in shipping boxes packed with ice and then stored in a -20 °C refrigerator until DNA extraction.

Plant genomes were extracted using liquid nitrogen homogenization and the DNeasy Plant Mini Kit (Qiagen, USA). Subsequently, the DNA’s quality and concentration were assessed by 1% agarose gel electrophoresis and Qubit 4 (Thermo Fisher Scientific, USA). Sequencing libraries were prepared from quality-checked total DNA samples by Novogene (Singapore), using theVAHTS Universal Plus DNA Library Prep Kit V4 (ND801, Vazyme). These libraries were then sequenced to produce 150-bp paired-end reads using a DNBSEQ-T7 platform (BGI, Shenzhen, China) by Novogene. Raw sequencing data were processed using Fastp (version 0.20.1) to remove adapter sequences and low-quality reads^20^. Additional genome sequence data were acquired from public databases, with accession numbers detailed in Supplementary Information Table S1.

### De novo assembly, annotation, and phylogenetic analysis of cp genomes

De novo assembly and annotation of cp genomes were preformed based on a method reported previously^15^. Cp genome assembly was conducted using GetOrganelle (version 1.7.7.1)^21^, with parameters set to −R 15, −k 21,45,65,85,105, and −F embplant_pt. Assembly validity was confirmed by mapping reads to the assembled sequence using Bowtie2 (version 2.4.4)^22^, with subsequent sorting and BAM file generation performed using Samtools (version 1.22.1)^23^. The cp genome assemblies were annotated using the online version of GeSeq^24^. The circular genome maps were generated using OGDRAW (version 1.3.1)^25^.

Phylogenetic analyses of cp genome sequences were conducted using HomBlocks (version 1.0)^26^ with default parameters to identify and align homologous regions, followed by trimming with trimAl (version 1.5. rev0) using the automated1 option. The resulting alignment was converted to PHYLIP format using EMBOSS Seqret. Maximum likelihood (ML) phylogenetic trees were inferred using PhyML (version 3.3.3)^27^ under the GTR model with a discrete gamma distribution (four rate categories) and empirical nucleotide frequencies, with branch support assessed using the approximate Bayes (aBayes) method. In addition, ML analyses with 1000 rapid bootstrap replicates were performed using RAxML (version 8.2.13) under the GTR+Γ model^28^. Phylogenetic trees were visualized using Dendroscope (version 3.8.10)^29^.

### Concatenated nuclear gene phylogeny (Read2Tree-based)

The concatenated phylogenetic analysis using Read2Tree (version 2.0.1)^14^ was also performed based on a method reported previously^15^. Marker genes for *Persea americana* and *Magnolia sinica* were obtained from the OMA browser (https://omabrowser.org/oma/export_markers). The OMA browser parameters were set with a minimum fraction of covered species at 0.8 and a maximum number of markers at –1. In total, 10,962 orthologous groups (OGs) were identified among the selected 2 species in the OMA browser. Read2Tree was run on paired-end whole-genome resequencing reads using the 2map step. OMA-derived marker gene sequences were supplied as a DNA reference. All other parameters were kept at their default values. To ensure the specificity of the phylogenetic trees, the 2 species obtained from the OMA browser were excluded from the resulting alignment files. Aligned data were trimmed using trimAl with the − automated1 option. ModelTest-NG (version 0.1.7) was used to identify the optimal evolutionary model^30^. Subsequently, Maximum Likelihood (ML) trees were generated using RAxML-NG (version 1.1) with 100 bootstrap replicates^31^. The visualization of the resulting trees was achieved using Dendroscope.

### Principal component analysis (PCA) for whole-genome sequence data

Prior to principal component analysis (PCA), nucleotide alignments for each orthogroup were extracted from the Read2Tree mapping outputs (05_ogs_map_*_dna/) using 16 ingroup taxa. Orthogroups represented in fewer than 50% of ingroup taxa or showing inconsistent alignment lengths among taxa were excluded. The retained orthogroup alignments were concatenated into a supermatrix using a custom Python script, resulting in a final alignment of 16 ingroup taxa spanning 16,454,499 sites. Missing taxa within retained orthogroups were represented as gap characters (-).

Variable sites were identified as positions exhibiting at least two nucleotide states (A, T, G, or C) among ingroup taxa. To reduce the influence of rare polymorphisms, only sites with a minor allele frequency (MAF) ≥ 0.05 were retained. Each polymorphic site was encoded as a binary variable (0 = major allele, 1 = minor allele), and ambiguous bases or gap-containing sites were imputed using the column-wise mean prior to analysis. After MAF filtering, 1,048,008 polymorphic sites were retained for PCA. PCA was conducted using the prcomp function in R (v4.5.3; R Core Team 2026) with mean-centering enabled and without variance scaling. The first three principal components (PC1–PC3) were visualised using ggplot2^32^, with taxa coloured according to species identity.

### Sparse non-negative matrix factorization (sNMF) analysis

Marker gene alignments for 16 *Cinnamomum* ingroup taxa were obtained using Read2Ttree, with *Persea americana* and *Magnolia sinica* as reference species, across 10,962 orthogroups. SNPs were extracted from per-orthogroup alignments using SNP-sites v2^33^ and concatenated across 10,918 orthogroups using bcftools^34^, yielding 1,031,664 SNPs across 16 taxa. Variants were then filtered by minor allele frequency (0.1 ≤ MAF ≤ 0.9) and randomly downsampled to 5% of sites, resulting in 35,220 SNPs for downstream analysis. Population structure was inferred using sNMF as implemented in the R package LEA^35^, with K = 2–8 ancestral components and 10 repetitions per K. The optimal number of ancestral components was determined by the cross-entropy criterion, which was minimized at K = 5. Ancestry proportion matrices (Q-matrices) were extracted for K = 3, 4, and 5 using the run with the lowest cross-entropy for each K, and visualized as stacked bar charts using ggplot2, with samples ordered according to their phylogenetic relationships.

### Use of Generative AI

ChatGPT (https://chatgpt.com/) was used for rewriting into more appropriate English and for English text proofreading. Afterward, the correctness of the revisions was confirmed by the authors.

### Ethics statement

Experimental research and field studies on plant material used in this study, including both cultivated and wild-collected samples, complied with relevant institutional, national, and international guidelines and legislation. *C. sieboldii*, *C. verum*, *C. insularimontanum*, and *C. × durifruticeticola* were obtained from the cultivated collection at the Tanegashima Division, Research Center for Medicinal Plant Resources, Kagoshima Prefecture, Japan, with the necessary institutional permission.

*C. yabunikkei* (Yakushima) and *C. sieboldii*_2 (main island of Kagoshima Prefecture) were collected from wild populations on private land with the permission of the landowners. Although

*C. sieboldii* is categorized as Near Threatened (NT) in the Kagoshima Prefecture Red Data Book, it is not designated as a legally protected species under Japan’s Act on Conservation of Endangered Species of Wild Fauna and Flora or under Kagoshima Prefecture’s conservation ordinances, and no special collection permit was therefore required beyond landowner consent. Both species were identified by Ei’ichi Iizasa. Voucher specimens of *C. yabunikkei* and *C. sieboldii*_2 were deposited at the Kagoshima University Museum (https://www.museum.kagoshima-u.ac.jp/) under accession no. KAG204593 and KAG204592, respectively.

## Data availability

The datasets generated and/or analyzed during the current study are available in the DNA Data Bank of Japan (DDBJ) Sequence Read Archive (DRA) repository (https://www.ddbj.nig.ac.jp/index.html), accession no. DRR1063129 to DRR1063134 and LC941049 to LC941054.

## Acknowledgements

We are grateful to Rinne Simizu, Yoshie Hamashima, Junichi Ebihara, Haruki Genozono, and Tomiyasu-Kashi (A Japanese confectionary shop) for providing fresh leaves of *C. yabunikkei* and *C. sieboldii*_2. We also thank Midori Wada and Yuka Kawaji for secretarial assistance. We would also like to thank Dr. Akihiro Asakawa, Dr. Katsuko Kajiya, Dr. Hisanori Tamaki, and Dr. Taiki Futagami of Kagoshima University and SAKURA HOS MEDICAL Co., Ltd. for their continuous support throughout this study.

## Author contributions

EI conceived and designed the project and revised the manuscript. SI assembled the sequences, analyzed the data, and wrote the manuscript. NA collected plant material and prepared photo plates. EI and SI acquired funding. All authors have read and approved the final manuscript.

---

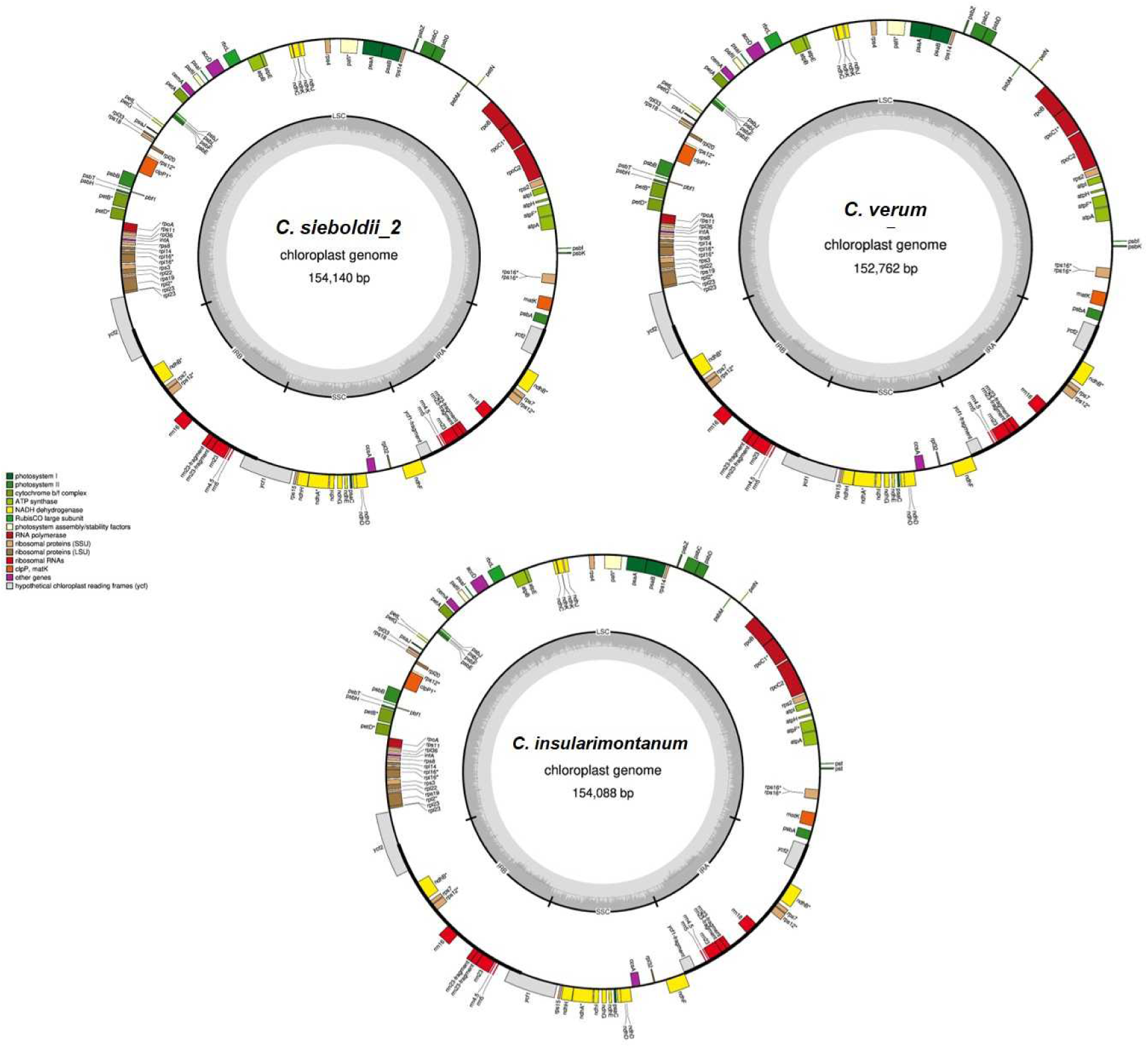

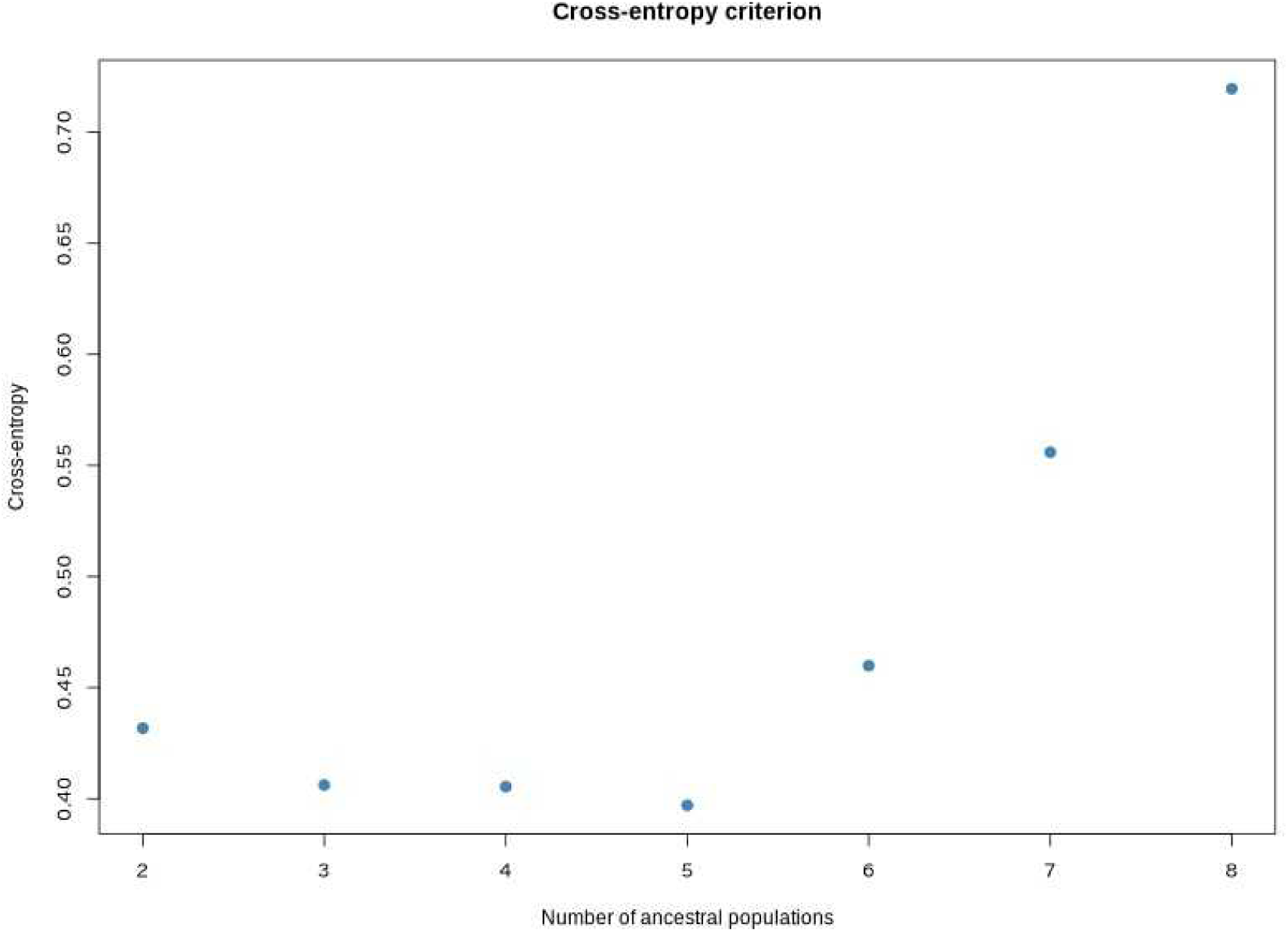

